# Genomic signatures of recent adaptation in a wild bumblebee

**DOI:** 10.1101/2021.05.27.445509

**Authors:** Thomas J. Colgan, Andres N. Arce, Richard J. Gill, Ana Ramos Rodrigues, Abdoulie Kanteh, Elizabeth J. Duncan, Li Li, Lars Chittka, Yannick Wurm

## Abstract

Behavioral experiments and analyses of observation records have shown that environmental changes threaten insect pollinators, creating risks for agriculture and ecosystem stability. Despite their importance, we know little about how wild insects or other animals can adapt in response to environmental pressures. To understand the genomic bases of adaptation in an ecologically important pollinator, we analyzed genomes of *Bombus terrestris* bumblebees collected across Great Britain. We reveal extensive genetic diversity within this population, and strong signatures of recent adaptation throughout the genome. More specifically, we find that selection recently affected key processes underpinning environmental interactions, including neurobiology, wing development, and response to xenobiotics. We also discover unusual features of the genome, including a 53-gene region that lacks genetic diversity in many bee species, and a horizontal gene transfer from a *Wolbachia* bacteria. The genetic diversity and gene flow we observe for this species could support its resilience to ongoing and future challenges. Overall, we provide important insight on the genetic health of an ecologically and economically important pollinator and reveal mechanisms by which it has recently adapted. The approach we used could help to understand how species differ in their adaptive potential, and to develop conservation strategies for those most at risk.

## Introduction

Behavioral experiments and analyses of observation records have shown that pesticide use, habitat fragmentation, emerging diseases and climatic change threaten insect pollinators including bees^1–4^. Despite the resulting risks for agricultural yields and for ecosystem stability, we know little about how wild insects may adapt to such environmental pressures. Similarly, we understand relatively little about how genomes are shaped in the wild^5^. If a species has adapted in response to a detrimental environmental pressure, then we should see changes in the alleles of genes involved^6,7^ (Fig. 1A). Analyzing genomes of many individuals can identify such changes and reveal the constraints and adaptive potential of species^8,9^. The resulting knowledge should support conservation efforts and practices^10^.

**Figure 1.**
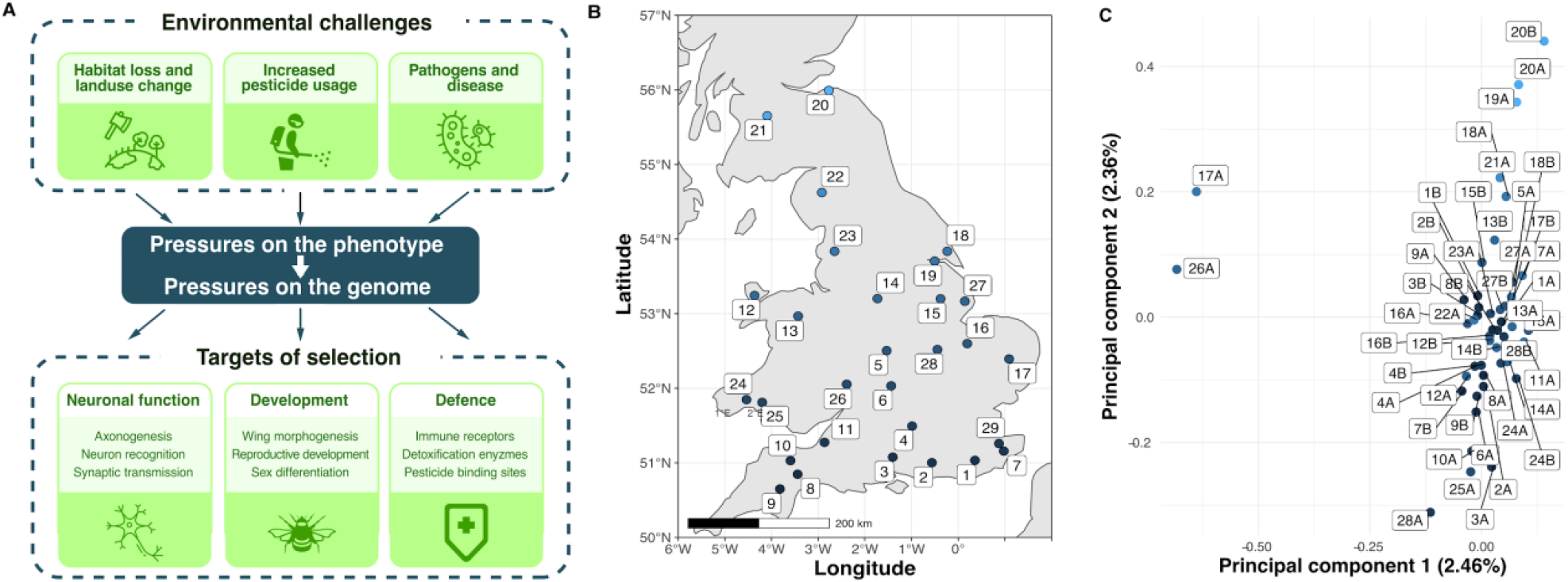
Environmental pressures affecting insect pollinators and population structure of wild-caught British *B. terrestris*. **(A)** Overview of key environmental selective pressures on wild bumblebee populations, and some of the biological pathways and processes expected to be under selection in response. **(B)** Twenty-eight collection sites across Great Britain, colored according to latitude. **(C)** Population structure of 46 males according to the first two principal components (PC1 and PC2). Each point refers to one male, with up to two males (A and B) per site, colored according to collection site latitude.

The annually reproductive bumblebee *Bombus terrestris* is ideal for understanding how coping with recent rates of environmental change is possible because, unlike many other pollinators, it has shown little evidence of population decline^1^. Furthermore, because male bumblebees are haploid, their genome sequences are unambiguous and intrinsically phased, providing more analytical power than the diploid genomes of female bumblebees and of many other insects. To understand which bumblebee genes and molecular processes underlie responses to recent selective pressures, we sequenced the genomes of male *B. terrestris* bumblebees from across Great Britain. We subsequently identified and characterized the genomic regions showing the strongest signs of recent adaptive evolution in this population. Our findings clarify the genetic health of this species and highlight genes and processes underpinning adaptation to recent challenges.

## Results

### Weak population substructure among *B. terrestris* in mainland Britain

We collected 46 unrelated male *B. terrestris* from across Great Britain and sequenced their genomes (411-fold total coverage; Fig. 1B; Supplementary Table 1). We found 1,227,312 single nucleotide polymorphisms (SNPs), with an average nucleotide diversity ***π*** of 1.51×10^−3^. To understand whether population structure constrains adaptation in this species, we performed identity-by-state and co-ancestry-based analyses. These analyses indicate that our dataset represents one population (Figs 1C and S1-2). Similarly, while the second principal component correlates with latitude (Pearson’s r=0.8, Figs 1C and S3), individual principal components explained at most 2.46% of genetic variation. Thus, despite the geographic heterogeneity of Great Britain, there is sufficient gene flow for these bees to be considered as one panmictic population, implying that new alleles have the potential to readily spread. The weak substructure of British *B. terrestris* is supported by studies using fewer markers^11–13^. Our result also indicates that no subset of our samples has the type of large-scale differentiation that could be expected from a cryptic subspecies.

### Selection is fine-tuning functional regions throughout the bumblebee genome

We used two approaches to identify signatures of recent selection in the genome. First, we identified large “hard” sweep regions, where selection on an allele can lead to haplotype fixation^14^. For this, we identified genomic segments longer than 100,000 nucleotides with significantly lower nucleotide diversity than the rest of the genome (z-score<−2σ). We found ninety such segments. Our second approach detected more localized signatures of selection, and “soft” sweeps, where two or more haplotypes are at high frequency. This can occur, for example, when strong selection on a new mutation occurs after a first allele reaches fixation, or when selection favors different alleles in different habitats^15^. For this, we determined for each SNP the metric |*nS*_L_| (“number of segregating sites by length”), a measure of haplotype homozygosity that is robust to variation in recombination and mutation rates; an |*nS*_L_| score greater than 2 is considered evidence of recent selection^16,17^. The 10,132 SNPs with the highest 1% of |*nS*_L_| scores have particularly strong signatures of recent selection (|*nS*_L_|≥2.56, *i.e.*, ≥3 standard deviations from the average; Fig. 2A; Supplementary Table 2) and are typically within 300 nucleotides of SNPs with low |*nS*_L_| scores. This indicates that recombination rapidly breaks down haplotypes in this species and that selection has acted in fine-tuned manners. The SNPs with high |*nS*_L_| scores are mostly in genic regions (79%; *p*<10^−15^) and, within these regions, are similarly represented in coding and non-coding sequence, indicating that selection recently acted on protein function and on gene regulation.

**Figure 2.**
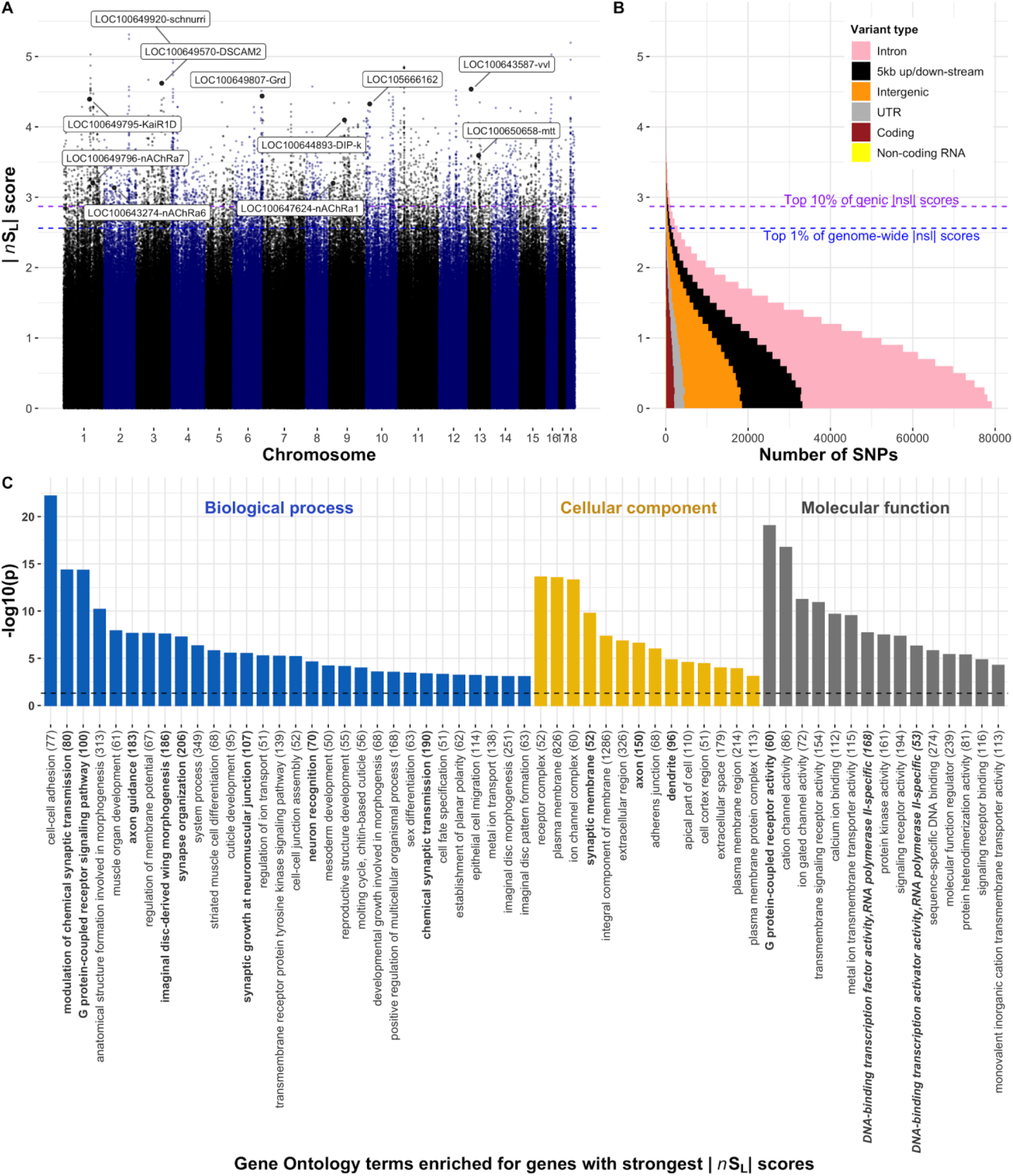
Genome-wide signatures of selective sweeps in British *Bombus terrestris* bumblebees. **(A)** |*nS*_L_| measures of selection for all SNPs in the bumblebee genome. Each dot represents one SNP; labelled dots represent the SNP with highest |*nS*_L_| score for genes of interest, including transcription factors, insecticide susceptibility genes and a *Wolbachia*-like gene, with high |*nS*_L_| scores. Labels indicate Flybase gene symbols when clear *Drosophila* orthology exists, otherwise, the NCBI gene symbol is provided. Blue and purple horizontal dashed lines respectively indicate the 1st percentile of overall |*nS*_L_| scores and 10th percentile of genic |*nS*_L_| scores. **(B)** Distributions of |*nS*_L_| scores show that most SNPs are in genic regions, and that most |*nS*_L_| scores are consistent with neutral or purifying rather than directional evolution as 96% of SNPs have |*nS*_L_| < 2. **(C)** Diverse Gene Ontology terms are enriched in genes with high |*nS*_L_| scores (−log10 transformed Bonferroni-adjusted *p* values). Terms associated with roles in neurology and transcription factor activity are respectively highlighted in bold and bold italics. The total number of annotated genes for each term is in parentheses.

To understand which types of biological functions were under the strongest recent selection pressures, we inspected annotations of genes with the strongest |*nS*_L_| scores and performed rank-based analyses of Gene Ontology and InterPro descriptions of all bumblebee genes (Bonferroni adjusted *p*<0.05; Figs 2C and S4; Supplementary Tables 3-4). The overviews of loci under recent selection, and the biological and molecular processes they affect, represent valuable resources for future phenotypic work on bees and on adaptation in natural insect populations (Figs 2C and S4; Supplementary Tables 3-4). Below, we highlight five particularly striking patterns regarding genes and regions with the strongest signatures of selective sweeps.

### Strong selection on transcription factors

Genes related to transcriptional regulation were overrepresented among genes under selection (Figs 2C and S4; Supplementary Tables 3-4). In particular, the gene with the strongest evidence of recent selection is the *B. terrestris* ortholog to the *schnurri* gene (|*nS*_L_|=5.14). In *Drosophila*, this transcription factor regulates embryonic patterning and wing patterning through the Decapentaplegic pathway^18^. Another transcription factor, the ortholog to the *ventral veins lacking* gene (*vvl*), has the 9th highest |*nS*_L_| score (|*nS*_L_|=4.54). In *Drosophila*, *vvl* is involved in steroid biosynthesis and embryonic brain development^19,20^, and intriguingly also interacts with the Decapentaplegic pathway to affect wing imaginal disc development^21^ and vein patterning^22^. These results, together with “wing morphogenesis” being the eighth most overrepresented Gene Ontology description among genes under selection, suggest that there was strong recent selection on wing structure. Such selection could be linked to recent changes to foraging or flight patterns^23,24^, because climatic changes modified the physical constraints of flying^25^, or perhaps in response to pathogens, such as the deformed wing virus, which can cause extensive wing abnormalities in infected individuals^26^.

### Strong selection acting on genes involved in bumblebee neurobiology

Neurological genes were overrepresented among genes with the highest |*nS*_L_| scores (Fig. 2C). In line with this, four of the 30 genes with the highest |*nS*_L_| scores have potential roles in neurotransmission (*gamma-aminobutyric acid receptor alpha-like*; |*nS*_L_|=4.44, *glutamate receptor ionotropic kainate 2*; |*nS*_L_|=4.39), axon guidance (*Down Syndrome cell adhesion molecule 2*; |*nS*_L_|=4.62), and memory formation (*neurotrimin*; |*nS*_L_|=4.1). Furthermore, selection on G protein-coupled receptor signaling in *B. terrestris* mirrors previous analyses on honeybees^27–29^. These receptor targets of hormones, pheromones and neurotransmitters are thus long-term targets of selection in social bees, potentially for roles responding to social or to environmental cues^30^. Selection on neurological genes could, for example, be linked to the need to improve complex cognitive and social behaviors of bumblebees^31^, for remembering increasingly complex foraging routes due to patchier habitats, or for neurological changes because of exposure to neurotoxins.

### Positive selection on a gene horizontally transferred from *Wolbachia*

We used similarity searches to understand the potential functions of uncharacterized genes among the genes with the twenty highest |*nS*_L_| scores. This showed that the gene with the 16th-strongest signature of selection (LOC105666162; |*nS*_L_|=4.33; Supplementary Table 2; Fig. 3) was horizontally transferred to *Bombus* from *Wolbachia*, a genus of bacterial endosymbionts that infect many invertebrates. Indeed, LOC105666162 has strong similarity to a gene in *Wolbachia* (BLASTP e-values<10^−16^) but not to most other insects (Supplementary Fig. S5; Supplementary Table 5; Supplementary Information). The data overwhelmingly suggest that this gene is integrated in the *Bombus* genome and not an artifact of potential contamination (Supplementary Information). Indeed, the same two orthologs flank LOC105666162 in *B. terrestris* and in *B. impatiens*^32^, and this gene is present in genomic sequences of 14 other *Bombus* species^33^, suggesting that horizontal transfer occurred 40 million years ago^34^. Additionally, sequencing depth across LOC105666162 is similar to the rest of the genome (paired t-test, t_df=95_=−0.73, *p*=0.47). Crucially, we find no other *Wolbachia*-related sequences in our samples, consistent with the absence of evidence that *Wolbachia* could infect *Bombus.* So, what might LOC105666162 do? Unfortunately, functional work on this gene and its *Wolbachia* homolog WD0147 are lacking, but we do have some clues. Horizontally transferred genes are often inactive, yet LOC105666162 is expressed in germlines and other tissues of both sexes and castes^35–37^, consistent with it being functional (Fig. 3; Supplementary Table 6). The *Wolbachia* homolog is expressed in infected *Drosophila* gonads^38^; based on its amino acid sequence and expression profiles, this gene is a strong candidate for driving mechanisms of cytoplasmic incompatibility^39^. One could speculate that LOC105666162 contributes to the lack of *Wolbachia* in bumblebees.

**Figure 3.**
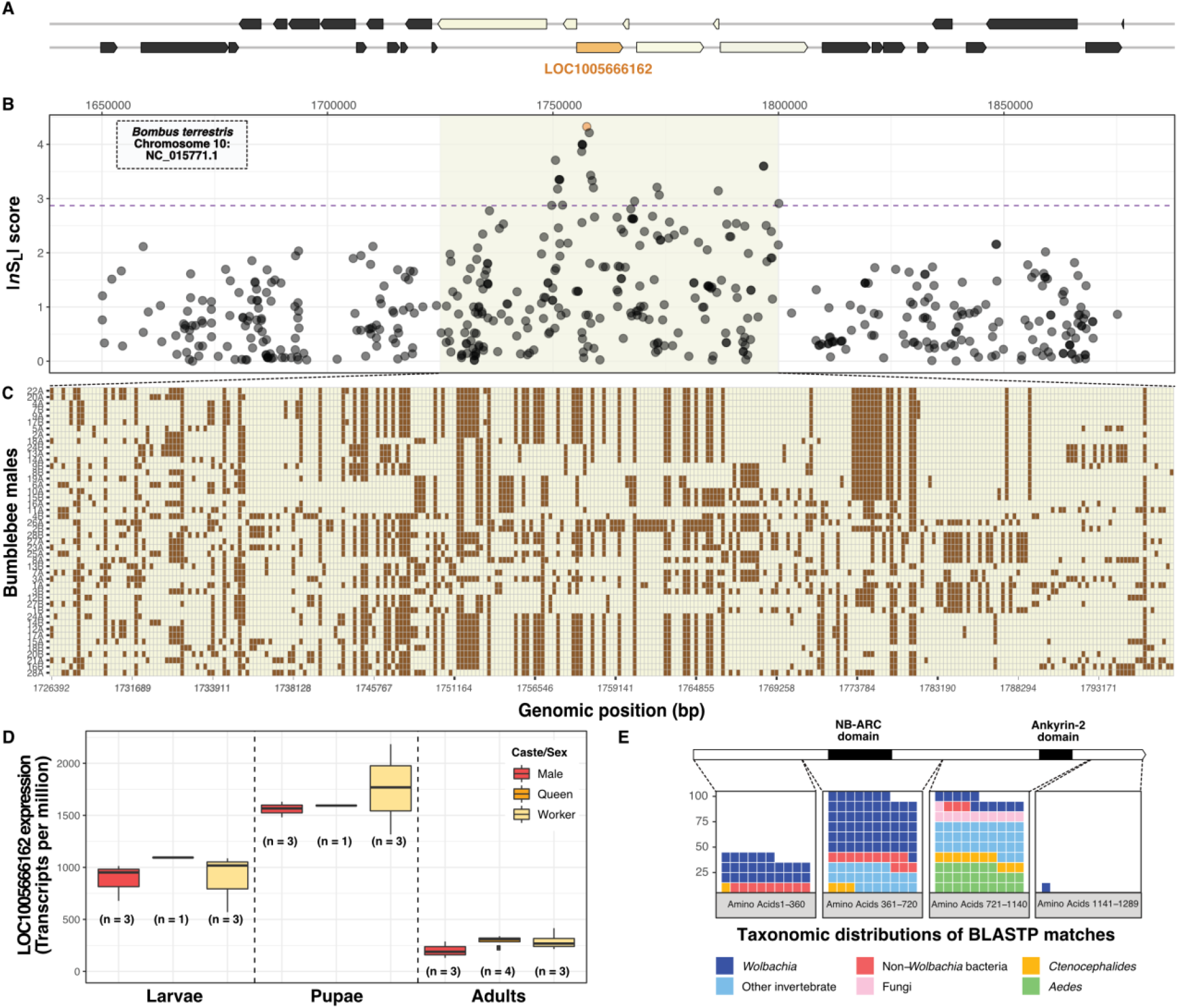
Positive selection acting on a *Wolbachia*-like ankyrin repeat domain-containing gene. **(A)** Genomic region surrounding LOC105666162 (orange). **(B)** |*nS*_L_| scores for each SNP in the region. LOC105666162 has the sixteenth strongest |*nS*_L_| score in the genome. **(C)** Genotypes for each of 46 male *B. terrestris* (rows) at each SNP (columns; chromosomal coordinates shown in the x axis). Colors indicate reference allele (beige) or alternative allele (brown). **(D)** LOC105666162 is expressed in male, queen and worker larvae, pupae and adults, indicating that it is likely functional across the life cycle of *B. terrestris*. **(E)** LOC105666162 gene structure includes NB-ARC and Ankyrin 2 domains. BLASTP searches with four sections of the gene, while excluding hits to *Bombus,* highlight strong similarity to *Wolbachia* and other bacteria, and limited similarity to other insects or fungi.

### An evolutionary conserved region of extremely low genetic diversity

A 200,000 nucleotide-long region stood out in our analysis because it contains only 56 SNPs and thus has 21-fold lower nucleotide diversity (**π**~7×10^−5^) than the genome-wide average (**π**=1.5×10^−3^; t_df=46_=90.4, *p*<10^−15^; Fig. 4). Furthermore, the low-diversity region has particularly high gene density (53 genes; z-score<−2σ) and represents a solid haplotype, with a population-derived recombination rate 298x lower (ρ=1.2×10^−4^) than the genome-wide average (ρ=0.035, t_df=1226700_=−460, *p*<10^−15^).

**Figure 4.**
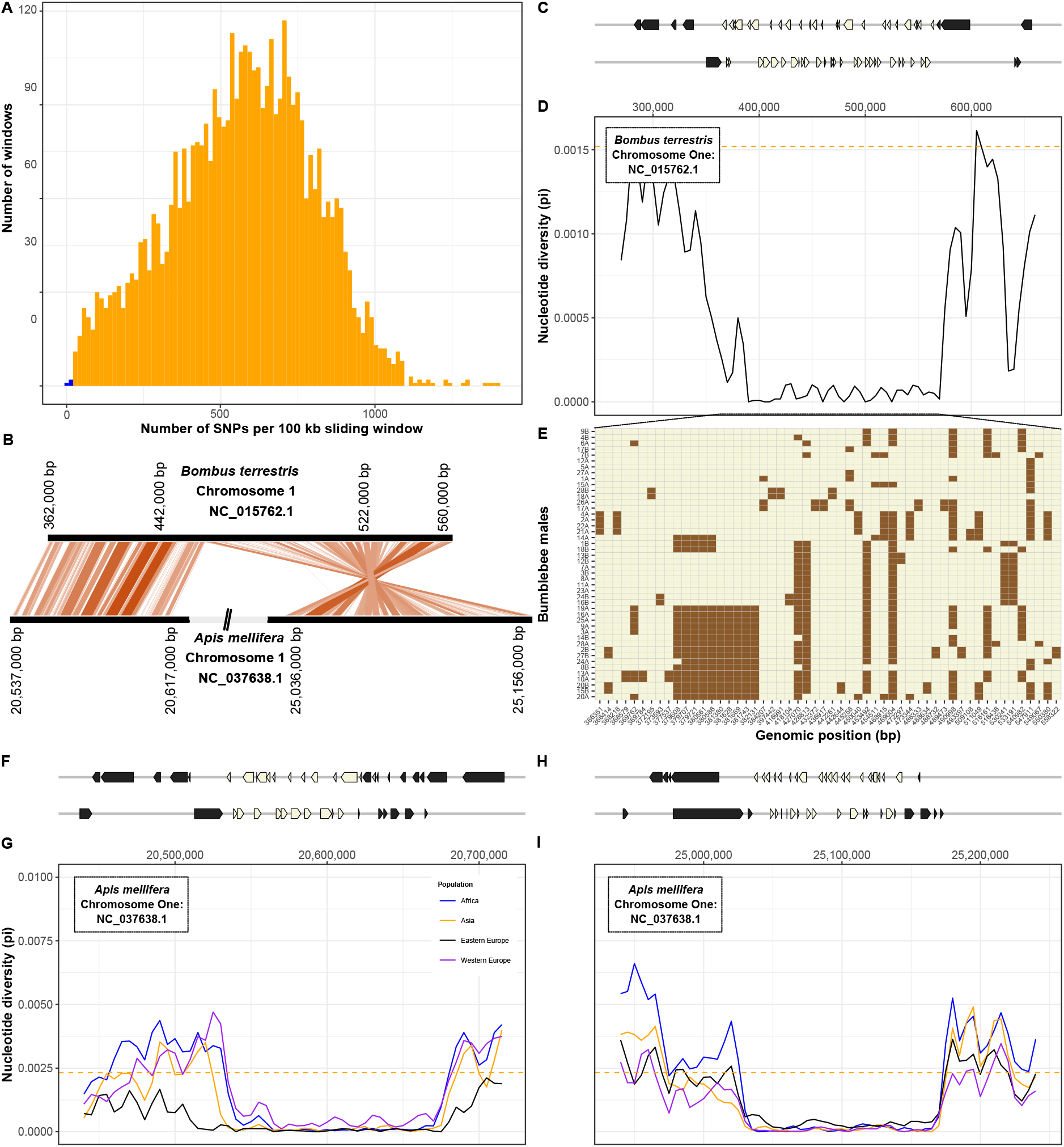
Conserved gene-rich region of low nucleotide diversity in bumblebee and honeybee. **(A)** Numbers of SNPs identified in 100 kb sliding windows across the bumblebee genome. The two windows with the lowest numbers of SNPs (in blue) are adjacent to each other on chromosome one. **(B)** Relative genomic positions of homologous regions of low diversity on chromosomes one of *Bombus terrestris* and *Apis mellifera*. **(C)** Genomic coordinates of 53 genes present in (beige) and flanking (grey) the region of low diversity. **(D)** Nucleotide diversity (**π**, calculated in 10 kb sliding windows) is low in this region in comparison to flanking regions and to the genome-wide mean (dashed line). **(E)** Genotypes for each of 46 *B. terrestris* males (rows) at each SNP (columns; chromosomal coordinate shown in the x axis). Colors indicate reference allele (beige) or alternative allele (brown). **(F-I)** In the honeybee *Apis mellifera,* homology with the region of low diversity in *B. terrestris* is split between two regions. For both regions, we show genomic positions of genes (F, H). In four populations, both regions have lower nucleotide diversity (**π**, calculated in 10 kb sliding windows) than flanking regions or the rest of the genome (dashed line; G, I).

To test whether the characteristics of this region are specific to *B. terrestris*, we identified orthologous regions in another bumblebee species *Bombus impatiens* and in the honeybee *Apis mellifera*. The orthologous region in *B. impatiens* contains 52 of the 53 genes and similarly has lower diversity (**π**~1.7×10^−4^) than other regions (genome-wide average **π**=1.2×10^−3^; t_df=18_=−22.8, *p*<10^−15^). Orthology to the honeybee is split between two regions separated by 4.4Mb, indicating that rearrangements have occurred since the common ancestor of bumblebees and honeybees existed 78 million year ago. For both regions, honeybee populations had at least 13-fold lower nucleotide diversity than the rest of the genome (region 1: t_df=19.328_=−98.9, *p*<10^−15^; region 2: t_df=15.8_=−65.2, *p*<10^−15^; Fig. 4). These patterns indicate that an intrinsic long-standing process is responsible for the low genetic diversity of these regions in bees. While the regions include genes that are likely under strong purifying selection (Supplementary Table 7), no particular gene annotation was overrepresented which could help interpretation. Unlike the rest of the genome, the lack of genetic diversity in this large region suggests that bees will have limited ability to adapt to selection pressures involving the genes it contains.

### Selection on potential insecticide susceptibility genes

Selection for resistance to neurotoxic pesticides can lead to changes in expression or sequence of target receptors or detoxification enzymes in insect pests^40^. Because bumblebees can be exposed to pesticides when foraging on crops, and given the extensively documented detrimental effects of pesticide exposure has on bumblebee health^3,4^, we preliminarily examined signatures of recent selection in known insecticide-response genes. Four target receptors of insecticides were among the 10% of genes with the highest |*nS*_L_| scores: three nicotinic acetylcholine receptor subunits which are targets of neonicotinoid insecticides (*nAChR1a*, *nAChR6a*, *nAChR7a*; |*nS*_L_|>3.12 for all), and *metabotropic glutamate receptor 2* (|*nS*_L_|=3.59), a target of the natural plant toxin L-canavanine^41^. Similarly, five cytochrome P450s and four carboxylesterases, which can contribute to detoxification, had strong signatures of selection (top 10% of genes with high |*nS*_L_| scores (|*nS*_L_|>2.89); Supplementary Table 8; Supplementary Fig. S6). This was also the case for 43 genes that are differentially expressed in bumblebees after exposure to neonicotinoid pesticides^37,42^ (Supplementary Table 9). Future research will help pinpoint the reasons for these patterns, and whether some of these changes may reduce susceptibility to toxins naturally present in pollen and nectar, or to synthetic pesticides.

## Discussion

Human-induced environmental changes add to long-standing ecological and evolutionary challenges faced by wild animals. Identifying potential causes of pollinator declines has to date relied on inferences from laboratory experiments or on correlative associations in the field. Our study takes an important step towards understanding the bases of resilience of an important pollinator species by uncovering signatures left in the organism’s genetic “blueprint” in response to selective pressures. The strong signatures of selection we find at loci throughout the *B. terrestris* genome are consistent with the view that insect pollinators face many different pressures. Environmental pressures likely contributed to recent changes we found to affect genes underlying physiology, neurology, and wing development.

Bumblebees also face intra- and interspecies pressures including, competition for food, habitat, and mates, and pressures from predators, pathogens and parasites. Large-scale gene expression and functional genomic datasets are only beginning to be produced for bumblebees^43^ and will be crucial for disentangling how the specific changes we observed affect phenotypes and fitness. Similarly, historical sampling of museum specimens could help characterize changes over time in morphology, population structure and allele frequencies.

The fine-tuning of adaptive responses in *B. terrestris* is highlighted by our finding strong signatures of selective sweeps within few nucleotides of neutrally evolving loci. Several characteristics of this species likely facilitate this. Crucially, the high recombination rate ^44^ and social lifestyle of *B. terrestris* mean that one queen can produce hundreds of haploid males, encompassing a broad diversity of allelic combinations. These males are fully exposed to the environment as they spend weeks foraging and trying to attract a mate^45^. Male bees are also subject to haploid selection, which should lead to faster adaptation than in diploid species^46^. Furthermore, the broad gene flow and large population size of *B. terrestris* enables the maintenance of large amounts of genetic diversity and the rapid spread of adaptive alleles. Future comparisons with sister species including those that are declining will clarify whether *B. terrestris* may have additionally harbored a generalist genetic toolkit further pre-disposing it to resilience.

Our approach shows that identifying recent signatures of selection can reveal how a wild pollinator has responded to the pressures it faces and can suggest its ability to respond to future pressures. Our work thus complements recent efforts in vertebrates and model systems. Future comparative genomic studies with other pollinators will improve our ability to disentangle why species differ in their resilience to recent environmental changes. Additionally, scaling up our approach will enable the creation of frameworks for predicting detailed responses to environmental challenges for entire ecological networks. Overall, while insect declines are worrying, we show how at least one common pollinator is adapting.

## Methods

### Bumblebee collection, DNA extraction and sequencing

In the summer of 2014 we collected up to two males from each of 28 sites, with each site being >20 km from the nearest neighboring site (Fig. 1B). Male *Bombus terrestris* (large earth or buff-tailed bumblebee) were caught using butterfly nets and transferred into individual 100 ml pots after morphological confirmation of sex and species. Pots were placed into a bag at 4-10°C. Within two hours, males were rapidly transferred to 2 ml cryotubes and then snap-frozen in liquid nitrogen. Subsequent storage was at −80°C.

From each bee, dissected tissue was homogenized in 200μl of phenol in a 2 ml screw-cap tube (Supplementary Table 1). Subsequently, DNA was extracted using phenol-chloroform followed by purification with the Sigma GenElute Mammalian Genomic DNA miniprep kit. DNA purity was initially assessed using a NanoDrop spectrophotometer (Thermo Fisher Scientific, UK) followed by quantification with a Qubit v3 fluorometer (Thermo Fisher Scientific, UK). DNA from each male was fragmented to ~550 bp using a Covaris M220 ultrasonicator and fragment size distribution assessed using a TapeStation 2200 (Agilent Technologies, UK). From each sample, we prepared an individually indexed Illumina TruSeq PCR-free DNA library, which was quantified using qPCR MasterMix (ABI Prism) and primer premix (Kapa Biosystems, UK). Libraries were pooled in equimolar concentrations and pairs of 125 bp sequences were produced on two lanes of Illumina HiSeq 2500 at Biomedical Research Centre Genomics, London, UK. Five samples were additionally sequenced on one lane of Illumina HiSeq 2500 at Oxford Genomics, Oxford, UK.

### Quality assessment and filtering of raw Illumina sequences

We obtained 616 million paired-end reads from the 51 samples we initially collected. Using bowtie2 (v.2.2.5;^47^) with the parameter ‘-X 1000’, we aligned raw reads to the *B. terrestris* reference genome (GCF_000214255.1;^−32^). The 422× cumulative genome coverage provided strong power to detect sites with nucleotide sequence polymorphism. Four males were removed from all biological analyses due to low coverage. A fifth male was only used for the analysis of contaminant sequences (Supplementary Information) because >58.1% of reads from this male lacked similarity to the reference genome. The mean mapped coverage for each of the remaining 46 samples was 11.8× (min: 7×; max: 26.7×). Quality of raw reads was assessed using FastQC (v.0.11.3; https://www.bioinformatics.babraham.ac.uk/projects/fastqc). Illumina adapters were detected and removed using Trimmomatic (v.0.33;^48^). Using Khmer (v.2.1.1;^49^) to first interleave pairs of reads, we removed sequences of low quality (where >25% of the read has a Phred quality score of strictly <20) using the fastx toolkit (v.0.0.14; http://hannonlab.cshl.edu/fastx_toolkit). We used Khmer to remove 31-mers present three or fewer times across the entire dataset, as they likely represent technical artefacts or particularly rare variants that we would be unable to analyze. Sequences shorter than 50 bp were removed using seqtk (v.1.0-r82-dirty; https://github.com/lh3/seqtk). The final cleaned dataset thus comprised 46 males with a mean coverage of 11.1× (min 6.7×; max 24.4×). This cleaned dataset provides sufficient power to genotype the majority of polymorphic sites because 96.2% of the genome had >1X coverage in each of the 46 males. Overall, ~99% of the reference genome had at least 20× coverage.

### Identification of polymorphic sites and genotyping of individuals

After mapping cleaned reads to the reference assembly using bowtie2, we called variants using freebayes (v.1.0.2-29-g41c1313;^50^) with the following parameters: --report-genotype-likelihood-max --use-mapping quality --genotype-qualities --use-best-n-alleles 4 --haplotype-length 0 --min-base-quality 3 --min-mapping-quality 1 --min-alternate-fraction 0.25 --min-coverage 1 --use-reference allele. We first removed the aforementioned five low-coverage individuals as they were each missing >10% of genotype calls, thus retaining data from 46 males. We then removed entire SNPs with low genotype quality scores (--minQ 20) and variants in collapsed repetitive regions (--max-mean-DP 100) using VCFtools (v.0.1.15;^51^). We removed sites that appeared to be heterozygous, which is impossible in haploids, and all sites with more than two alleles as they also likely represent collapsed regions in the reference genome. To further reduce dataset complexity, we used --remove-indels to only consider single nucleotide polymorphisms (SNPs). We calculated allele frequencies and retained genotypes only where the rare allele was present in at least two males. Finally, we only considered those SNPs in regions of the genome that are mapped to the 18 linkage groups (representative of chromosomes). Mean nucleotide diversity ***π*** was calculated using 10 kb sliding windows with 5 kb overlap using PopGenome (v.2.2.4;^52^).

### Assessment population structure

We investigated among collected bumblebees by performing identity-by-state (IBS) analyses on a pruned set of SNPs generated by SNPRelate (v.1.8.0;^53^) using parameters that are similar to those previously used for *Drosophila*^54^ (--ld-threshold=0.2 --slide.max.n=500). We further investigated population structure using three approaches with unpruned SNPs: principal component analysis using SNPRelate, ADMIXTURE (v.1.3.0;^55^) with *K=*1-40 using cross-validation (--cv) as a measure to identify the best *K* value, and the linkage-aware approach fineSTRUCTURE (v.0.1.0;^56^).

### Evidence of recent selective sweeps

First, we identified regions of the genome with particularly low nucleotide diversity, indicative of "hard" sweeps. Second, to identify potential “soft” selective sweeps, we calculated *nS*_L_^16^ for all high confidence SNPs using selscan (v.1.1.0b;^17^). This metric is a measure of extended haplotype homozygosity. We normalized all *nS*_L_ scores against the empirical genome-wide distribution using selscan “norm”, using default settings. We used the top 1% (|*nS*_L_|≥2.56) absolute score of the *nS*_L_ metric (|*nS*_L_|) for all downstream analyses because (|*nS*_L_|≥2) indicates a selective sweep. Normalized |*nS*_L_| scores per gene, as well as NCBI RefSeq gene symbol and description, are provided in Supplementary Table 2.

### Data availability

Sequencing data are available from NCBI BioProject (PRJNA628944).

## Acknowledgements

We thank Richard Nichols, Rodrigo Pracana, Eckart Stolle, Carlos Martinez Ruiz and Gino Brignoli for advice on analysis and wording. We thank Monika Struebig and Christopher Durrant for help with molecular work. We also thank BRC Genomics, London, and Oxford Genomics, Oxford, UK, for the sequencing of genomic libraries. This work was supported by the Biotechnology and Biological Sciences Research Council (grant BB/K004204/1), the Natural Environment Research Council (grants NE/L00626X/1 and NE/L00755X/1), NERC EOS Cloud and QMUL Research‐IT and MidPlus computational facilities (The Engineering and Physical Sciences Research Council; grant EP/K000128/1).

## Author Contributions

YW, RJG, LC, TJC and ANA conceived, designed the project and coordinated collections. ARR helped collect wild bumblebees. LL performed dissections. TJC performed most genome-level analyses, with assistance from YW and AK. EJD helped with interpretation of results. TJC and YW drafted the manuscript. All authors wrote and revised the final manuscript.

