## Supplementary Information for "Genomic signatures of recent adaptation in a wild bumblebee"

**This PDF file includes:**

**Supplementary Materials and Methods** **2**

Computational infrastructure 2
Confirmation of species identity using morphology and mitochondrial sequence 2
Inferring potential effects of SNPs in coding sequences 2
Assessment of population structure 3
Assessment of non-*Bombus* sequences within bumblebee reads 3
Evidence for horizontal gene transfer event in *Bombus* genomes 4
Phylogenetic analysis of horizontal gene transfer 5
Association of copy number variation with recent selective sweeps 6
Presence of insecticide-response genes within recent sweeps 7
Limited signatures of recent positive selection on canonical immune genes 7
Comparative genomic analysis for an evolutionarily conserved region of low genetic diversity 7
Assessment of macroevolutionary processes acting on genes in regions of low diversity 9
Functional annotation of gene-rich region of low genetic diversity in *B. terrestris* 9
High prevalence of the endosymbiotic bacteria *Arsenophonus* in one bumblebee male 10
InterPro domain and Gene Ontology term enrichment analysis 11

**Supplementary Figures** **12**

Supp. Figure S1. Identity-by-state analysis for wild-caught bumblebees. 12
Supp. Figure S2. Weak population substructure within British *B. terrestris.* 13
Supp. Figure S3. Correlation between genotype frequencies and latitude. 14
Supp. Figure S4. Enriched functional domains in bumblebee genes under recent selection. 15
Supp. Figure S5. Phylogeny of LOC105666162, the gene horizontally transferred from *Wolbachia* into *Bombus* and similar sequences from other organisms. 16
Supp. Figure S6. Putative pesticide-response genes under selection in the bumblebee genome. 17

**Legends for Supplementary Tables** **18**

**Supplementary References** **20**

**Other supplementary materials for this manuscript include the following:**

Supplementary Tables 1 to 10

### Supplementary Materials and Methods

#### Computational infrastructure

Data were processed and analyses performed using snakemake scripts (v.3.2.1;[1]), bash commands and R scripts. SNPs were analyzed and summary data were visualized using R (v.3.5.1; R Core team (2018)). Genome mappings were visualized using the Integrative Genomics Viewer (v.2.5.3;[2]) and Ensembl[3]; BLAST analyses of individual genes and regions were performed using SequenceServer[4]. Computation was performed on Apocrita Midplus running CentOS 6 (https://docs.hpc.qmul.ac.uk).

#### Confirmation of species identity using morphology and mitochondrial sequence

Species and sex of individuals were identified based on color banding and morphology during field collection. We also compared raw sequence data for each male against *Bombus* mitochondrial cytochrome *c* oxidase 1 (CO-I) nucleotide sequences obtained from the Barcode Of Life Data (BOLD) database (Bold Systems v.4). We reduced redundancy by clustering all downloaded *Bombus* CO-I sequences using CD-HIT (v.4.6.8;[5]) and collapsing sequences that shared >97% sequence similarity into consensus sequences. We created a custom BLAST database with these consensuses using Magic-BLAST (v.1.4.0;[6]). We compared sequences from each male against this database, retaining only reads that shared 100% sequence similarity with sequences present in the database. We then counted the number of reads aligning to consensus sequences originating from different *Bombus* species present in the database. For all individuals, the highest number of matches were against BOLD sequence BEEEE113-15, which originates from *B. terrestris audax*, the subspecies found in Great Britain.

#### Inferring potential effects of SNPs in coding sequences

We used SnpEff (v.4.3;[7]) to predict the functional effects of alternate alleles for all 1,227,312 high confidence SNPs. Among the coding variants identified in *B. terrestris*, 10,429 (0.8%) are nonsynonymous. A total of 348 SNPs, across 302 protein-coding genes, were putatively high impact, with the following effects: "loss of start codon" (n=15), "gain of stop codon" (n=123), "loss of stop codon" (n=49), "splice acceptor variant and intron variant" (n=59), "splice donor variant and intron variant" (n=92), "stop gained and splice region variant" (n=6), and "stop lost and splice region variant" (n=4). While most high impact alleles were present only at low frequency, 18 were present as alternate alleles in more than 75% of individuals, indicating that the reference genome includes alleles that are rare in the British population and could be detrimental.

##### Assessment of population structure

We assessed the population structure of *B. terrestris* in mainland Britain using four approaches. First, identity-by-state analyses found that the proportion of alleles shared amongst individuals fit the expectation that they are from a single population (Supplementary Fig. S1). Second, fineSTRUCTURE clustered all individuals into a single population based on shared ancestral haplotypes. Third, ADMIXTURE analysis also indicates that our dataset represents one population (*k*=1; Supplementary Fig. S2). Finally, a principal component analysis after pruning SNPs in approximate linkage (n=1,015,887 SNPs; Fig. 2B), identified weak population structure with the strongest principal components (PCs) each explaining <2.5% of genetic variation. The first PC separated two males from all others, while the second PC correlated with latitude (Pearson's r=0.8, Fig.2B; Supplementary Fig. S3).

##### Assessment of non-*Bombus* sequences within bumblebee reads

To determine whether any microbial sequences are present in our samples, we first aligned raw reads against the *B. terrestris* reference genome using bowtie2. Unmapped reads were output into a separate directory and subsequently taxonomically classified using Kraken[8] with a database of bacterial, fungal and viral genomes generated using scripts provided by Kraken (downloaded January 19th, 2018). One bumblebee male stood out in that only 43% of filtered reads aligned to the reference genome. The majority of the remaining reads (82%) matched *Arsenophonus nasoniae* (GCA_000429565.1), an endosymbiont of *Nasonia* wasps that distorts sex ratios. This wild-caught bumblebee thus was highly infected by *Arsenophonus.*

We performed an additional analysis to test whether our males contained sequences from the intracellular symbiont *Wolbachia.* First, we downloaded all 43 *Wolbachia* genome assemblies available from the NCBI RefSeq database (as of January 25th, 2017). We then used bowtie2 to align raw reads for each bumblebee against a combined database containing the *Wolbachia* genomes and the bumblebee reference genome. Fewer than 1% (mean 5,686 reads) of unmapped reads aligned to any of the *Wolbachia* genomes. Manual inspection of these alignments indicated that these were spurious matches that had low mapping quality scores. This suggests that no *Wolbachia* are present in our sampled bumblebees.

**Evidence for horizontal gene transfer event in *Bombus* genomes**

In our analysis, LOC105666162 was the gene with the 16th highest $|nS_L|$ score and has high similarity to a gene in the bacterial endosymbiont *Wolbachia*. To determine sequence coverage at LOC105666162, we calculated genome-wide read depth per base using samtools depth (v.1.2;[9]). We compared mean coverage across the gene with flanking regions on 10 kb of either side of the gene, as well as against the genome-wide average. For calculating the genome-wide average, we removed extreme counts for bases from highly repetitive regions (depth>100).

To determine whether LOC105666162 exists in other *Bombus* species, we combined two approaches. First, a simple comparison with the *B. impatiens* genome indicates it includes a clear ortholog to LOC105666162, which is flanked by orthologs to the same two genes as in *B. terrestris.* Genome assemblies are unpublished for 12 other *Bombus* species, thus we downloaded their raw sequence reads from SRA[10] and aligned them against *Bombus terrestris* chromosome 10 (NC_015771.1) using bowtie2. Samtools depth was used to calculate the percentage of mapped reads. BAM files were indexed using samtools and visually inspected using IGV. While sequence divergence was obvious, we found no particular pattern of lower coverage, unpaired reads or mismatched read pairs. This confirms that one-to-one orthologs of the focal gene and its immediate neighbors are present in synteny in the other bumblebee genomes. To obtain a consensus sequence from each species, we called variants using freebayes. We retained variants called in all species (*i.e.,* no missing sites) with a minimum quality score of 20. We used bcftools 'consensus' (v.1.3;[11]) on species-specific VCFs to generate individual FASTA files for the coding sequences for the predicted orthologs of LOC105666162 and calculated *dN/dS* ratios between each species (range: 0.35-1.15).

In the Ensembl gene annotation, LOC105666162 codes for two protein isoforms of length 1289 and 1154aa, respectively. The shorter isoform has a later translation start site but otherwise overlaps

100% with the larger isoform. The longer isoform contains two types of functional domains, an NB-ARC domain (pfam00931; amino acid (aa) positions 385-566) and at least one ankyrin repeat domain (pfam12796; aa positions 986-1080). The predicted protein lacks a signal peptide or transmembrane domain suggesting intracellular function. The second section of the protein sequence, which contains the NB-ARC domain, shares sequence similarity with bacterial sequences, primarily *Wolbachia* (Fig. 3, Supplementary Table 5). The NB-ARC domain has similarity to sequences in genomes of a small number of eukaryotes (22 proteins coded for by 19 unique genes), which were not consistent with shared ancestry (Supplementary Table 5). For two of these eukaryotic species for which genome assemblies are available, the pattern differs substantially from what we see in *Bombus*, in that flanking genes also have high sequence similarity to *Wolbachia* genomes. This suggests that in those species, a larger segment of the *Wolbachia* genome recently integrated into the host genome or that the region represents contamination from an extant symbiont. For example, both neighboring genes for the ant *Vollenhovia emeryi* code for proteins with next top matches against proteins from *Wolbachia* species (percentage identity >98%, query coverage >99%). For the other species, *Nasonia*, sequence similarity to *Wolbachia* sequence was lower (percentage similarity >32.5%, query coverage >95). Our focal bumblebee gene (LOC105666162) is flanked by two genes that are highly conserved across eukaryotes (*CXXC motif domain-containing zinc-binding*: LOC100650449; *phosphatidate phosphatase LPIN3*: LOC100649067). These flanking genes have high nucleotide sequence similarity to their orthologs in honeybee (>=84.3% identity BLASTN against NCBI nt database; e-values <$10^{-131}$). While the two *Apis* orthologs are adjacent on chromosome ten, no gene in any part of the honeybee genome shares sequence similarity with LOC105666162.

**Phylogenetic analysis of horizontal gene transfer**

We performed a phylogenetic reconstruction of sequences with similarity to LOC105666162 (Supplementary Fig. S5). The resulting phylogenetic reconstruction of the identified sequences does not resemble the tree of 15 taxa which carry homologs. This pattern suggests that some of the sequences are mislabelled – perhaps due to contamination or assembly artifacts, or that horizontal gene transfer occurred independently in multiple genera, or both. When orthologs to LOC105666162 were flanked by genes with unambiguously eukaryotic ancestry, we found no conservation of synteny, further indicating that independent acquisitions may have occurred.

**Association of copy number variation with recent selective sweeps**

We identified copy number variation (CNV), *i.e.,* duplications and deletions within our dataset. First, we identified putative CNV sites using lumpySV (v.0.2.12;[12]). We retained all potential deletions but removed potentially duplicated sites <500 bp. For all putative CNVs, we removed regions with an excess of ambiguous bases in the reference genome (>10%), identified using seqtk comp, and regions with an excess of multiple mapping reads (mapQ=0). Such regions may represent highly repetitive or low complexity sites. Second, we calculated read depth across putative CNV sites. In brief, we aligned filtered reads against the *B. terrestris* reference genome (GCF_000214255.1) using speedseq (v.0.1.0;[13]). For each putative CNV, read depth was calculated for each individual using samtools depth. These were compared against the median read depths of 845 randomly chosen sites of comparative sizes using bedtools2 shuffleBed (v2.24.0-2-gb3504ce;[14]). This was performed independently for duplicated and for deleted regions. Putative duplications were identified as sites with a median read depth >2 standard deviations higher than the mean of the randomly chosen regions.

We identified 135 putative duplicated genomic regions within which at least one male had read depth at least three standard deviations than the median value of genome-wide read depth. Twenty one of the regions had elevated read depth in all males suggesting that these regions are duplicated throughout the British population. Among the 135 regions, 81 overlapped with genes. This high proportion (60%) suggests that many of the copy number variants are or were adaptive. Twenty of the 135 regions overlapped with more than one gene, suggesting that these copy number variants affect function or expression of multiple genes. We also found copy number variation for two cytochrome P450 genes, which are genes important for detoxification, although neither gene had a high $|nS_L|$ score nor was differentially expressed in response to pesticides (Supplementary Table 10).

#### Presence of insecticide-response genes within recent sweeps

*B. terrestris* homologs of *D. melanogaster* genes annotated with a "response to insecticide" (GO:0017085) GO term were obtained from Ensembl Metazoa BioMart. We additionally examined major detoxification enzyme families including cytochrome P450s, glutathione-S-transferases and carboxylesterases. To ensure confidence in polymorphic sites with signatures of selection ($|nS_L|>2.56$), manual inspections were performed for 500 SNPs of interest using VCF and individual BAM files using IGV. For each SNP, we examined the genomic position, genotype call, and 200 bp surrounding each SNP.

#### Limited signatures of recent positive selection on canonical immune genes

Pathogens can place enormous selective pressures on hosts to evolve and adapt defenses[15]. One such defense is the immune system, a key barrier to infection and establishment of disease[16]. Given the pressures placed on hosts by pathogens, certain genes involved in the host immune system are expected to be under recurrent positive selection[17]. In our study, while some canonical immune genes (n=12 of 179;[18]) were in the top 10% of genes with the highest $|nS_L|$ scores ($|nS_L|$ score>2.87), we did not identify significant enrichment of immune-associated GO terms (Bonferroni adjusted $p>0.05$) in genes with evidence of recent selection. The absence of pattern may be at least in part due to a paucity of molecular data regarding bumblebee immunity.

#### Comparative genomic analysis for an evolutionarily conserved region of low genetic diversity

An approximately 200,000 nucleotide region of chromosome one containing 53 genes stood out as having particularly low nucleotide diversity (Fig. 4): ~21-fold (mean $\boldsymbol{\pi}=7.3\text{x}10^{-5}$) lower than the genome-wide mean ($\boldsymbol{\pi}=1.5\ \text{x}10^{-3}$; $t_{df=46}=90.4$, $p<10^{-15}$). In line with this, the mean distance between consecutive SNPs was higher (3,700 bp) than the genome-wide average of 177 bp ($t_{df=55}=5.7$, $p<10^{-6}$), and the population-estimated recombination rate in this region is 298 times lower (mean rho=1.2<$10^{-4}$) than the genome mean (rho=0.035, $t_{df=1226700}=-460.53$, $p<10^{-15}$). The entire region contains 56 polymorphic sites of which approximately half (n=26) are within 5,000 bp upstream of genes. In contrast to genome-wide SNPs where 56.9% of polymorphic sites are located in intronic regions, this region only has one intronic variant (<1% of region-specific SNPs). Similarly, the region contains no intergenic SNPs (genome-wide mean is 99 intergenic SNPs per 100,000 bp $t_{df=2155}$ =-

24.5, $p<10^{-16}$). The greatest distance (>19 kb) between consecutive SNPs was between chromosomal positions 397,442 and 416,991, which contains three complete genes (LOC10064609: *cilia- and flagella-associated protein 91*; LOC100645973: *F-box only protein 28*; LOC100645666: *importin-7*). For twenty six genes in the region, there were no SNPs within 5,000 bp. While the region is not enriched for any particular Gene Ontology term, the region contains genes annotated with potential roles in ubiquitination, ribosomal function, neurotransmission, chemoreception and immunity.

Reductions in nucleotide diversity can be due to strong positive or purifying selection. Comparative analyses with other species can shed light on this. We identified orthologs of the region's 53 *B. terrestris* genes in *B. impatiens* and *Apis mellifera*. Synteny with *B. impatiens* was high with 52 of 53 genes located in a region of 196.5 kb (scaffold NT_176796.1: positions 470,040 - 669,574). We identified clear orthologs in honeybee for 46 genes, but these were split between one block of 16 orthologues and a second block of 30 orthologues 4.4 Mb downstream of the first synteny block. There was higher similarity (88.75%) of orthologous genes in the region between *B. terrestris and A. mellifera* than in the rest of the genome (87.5%) ($t_{df=44.43}$=-2.02, $p<10^{-3}$), suggesting that perhaps purifying selection plays a strong role in this region. In line with this, using 22 genes identified by OrthoDB as single copy orthologs across five bee species, 19 were under purifying selection ($dN/dS<1$ using the branch-site model).

To further understand the evolutionary history of the highly conserved genomic region, we calculated nucleotide diversity across the honeybee reference genome using population genomic data derived from 39 workers collected across three continents and representing four *A. mellifera* subspecies[19]. We similarly obtained whole genome sequence data from 10 male *Bombus impatiens*[20]. For each species, we aligned reads to their respective genome assemblies using bowtie2 (GCF_003254395.2 (Amel_HAv3.1;[21]); GCF_000188095.1 (BIMP_2.0;[22])).

In the red fire ant *S. invicta*, we identified 47 orthologues to the 53 genes in the *Bombus terrestris* region. However, the 47 *S. invicta* genes were split across 17 genomic scaffolds, showing that the region is not syntenic at this evolutionary distance.

#### Assessment of macroevolutionary processes acting on genes in regions of low diversity

We investigated evidence of long-standing signatures of positive or purifying selection acting on the 53 genes present in the low-diversity region. We aimed to test whether they were single copy orthologs and whether such genes had signatures of accelerated rates of positive or purifying selection that might be consistent with recurrent selection. For this approach, we obtained single copy orthologs from OrthoDB (v.9;[23]) for five bee species: *B. terrestris, B. impatiens*, *A. mellifera*, *Megachile rotundata* and *Habropoda laboriosa*. We performed a PRANK alignment (v.170427; parameters: -showtree -codon +F;[24]) on each set of protein-coding single copy orthologs. We removed gap-rich regions using Gblocks (v.0.91b; parameters: type=codon, minimum length of block=6, no gaps allowed;[25]). We identified signatures of positive selection using the branch-site model in codeML (PAML v. 4.8;[26]).

#### Functional annotation of gene-rich region of low genetic diversity in *B. terrestris*

Among the 53 *B. terrestris* genes in the region of low genetic diversity, 52 were expressed in all *B. terrestris* life cycle stages, including larvae and pupae, as well as both sexes using data previously generated by Harrison et al.[27] and Lewis et al.[28]. There was no general difference in sex-biased expression between genes in the region of interest and the rest of the genome. This suggests that the genes are subject to haploid selection, but that it is likely overall at a similar level as other genomic regions[29]. However, 15 genes were more highly expressed in adult males than workers (Benjamini-Hochberg corrected $p<0.05$). Five of these genes were single copy orthologs with the majority of coding sites (>85%) categorized by PAML as being under purifying selection within *Bombus* and in the other bee species examined. This was similar to the patterns in the rest of the genome ($t_{df=21.074}=0.66261$, $p=0.5148$).

The cause for the conserved low diversity in the two groups of social bees is unclear. Low recombination, positive selection and background selection can all result in regions of low diversity, and male haploidy is thought to increase the efficacy of selection. Perhaps a combination of processes is responsible for the particular characteristics of this region. Alternatively, it is tempting to speculate that the entire region may have recently introgressed from a related species. However, it is difficult to imagine this occurring independently in multiple species unless the regions also somehow contain selfish meiotic distorters. Perhaps the simplest explanation is that strong haploid selection contributes to the creation of a runaway process – a genomic equivalent of a black hole. Appropriate functional genomics work will help to reveal the processes taking place. The 53 genes in the region are involved in important cellular processes, such as ubiquitination (e.g. *ubiquitin-like-conjugating enzyme ATG3*; *E3 ubiquitin-protein ligase BRE1A*; *F-box only protein 28*; *NEDD8-conjugating enzyme Ubc12*) and ribosomal function (*60S ribosomal protein L21*; *oligoribonuclease*) but the region also contains genes with putative roles in chemoreception (*ejaculatory bulb-specific protein 3*), neurotransmission (*G-protein coupled receptor 139*; *KV channel-interacting protein 4*) and immunity (*FAS-associated factor 1*) which may be expected to be polymorphic due to strong selection pressures placed on the genes through environmental interactions. Furthermore, this region also contains genes coding for long non-coding RNA which have intriguingly high levels of nucleotide similarity conserved between *B. terrestris* and *A. mellifera*.

An interesting candidate for future research is the *pi3k* gene, which for both *B. terrestris and A. mellifera* was the most polymorphic gene in this conserved region of low nucleotide diversity. The gene is homologous to *vacuolar protein sorting 15*, a gene in *Drosophila melanogaster* that functions in the VSP34/Class III PI3-kinase complex and the production of phosphatidylinositol 3-phosphate, which is essential for regulation of autophagy and endocytosis[30,31]. Through its role in the PI3-kinase complex, the gene has also been implicated in neuron remodeling[30] and antibacterial immune regulation[31]. Future research will be required to elucidate the molecular function of these 53 genes, as well as decipher the evolutionary history and consequences of this linked haplotype of low nucleotide diversity in these social insects.

##### High prevalence of the endosymbiotic bacteria *Arsenophonus* in one bumblebee male

A benefit of whole-genome sequencing is that there is the potential to sequence organisms resident within or on the host organism at time of extraction. While other pathogenic organisms may have been present in our sampled bees, for one bumblebee male, taxonomic classification of non-*Bombus* aligned reads identified the most abundant taxonomic group as an *Arsenophonus* endosymbiont. *Arsenophonus* are a clade of intracellular symbiotic bacteria found in a diverse range of insect hosts[32] and have been detected within the digestive tract of wild and commercial bumblebees in

Great Britain[33]. *Arsenophonus* can be transmitted horizontally among honeybees[34], and diseased honeybee colonies have high *Arsenophonus* prevalence[35], raising concerns about a potential role in host pathology. Further research will be required to understand how *Arsenophonus* may affect fitness or physiology of beneficial pollinator species.

**InterPro domain and Gene Ontology term enrichment analysis**

We downloaded InterPro domains annotations for *B. terrestris* genes from Ensembl Metazoa BioMart. We ranked all genes with $|nS_L|$ scores from highest to lowest. For each InterPro domain present at least 20 times, we tested whether it was overrepresented among genes with high $|nS_L|$ using a Wilcoxon rank-based test.

Because little functional information exists for *B. terrestris,* we used Gene Ontology annotations from the best *D. melanogaster* ortholog for each *B. terrestris* gene, as provided by Ensembl Metazoa BioMart. We tested for GO term enrichment by applying $|nS_L|$-rank based Kolmogorov–Smirnov tests using topGO (v.2.26.0;[36]) with the "weight01" algorithm (nodeSize=50). We corrected for multiple testing using the Bonferroni method (adjusted $p<0.05$).

### Supplementary Figures

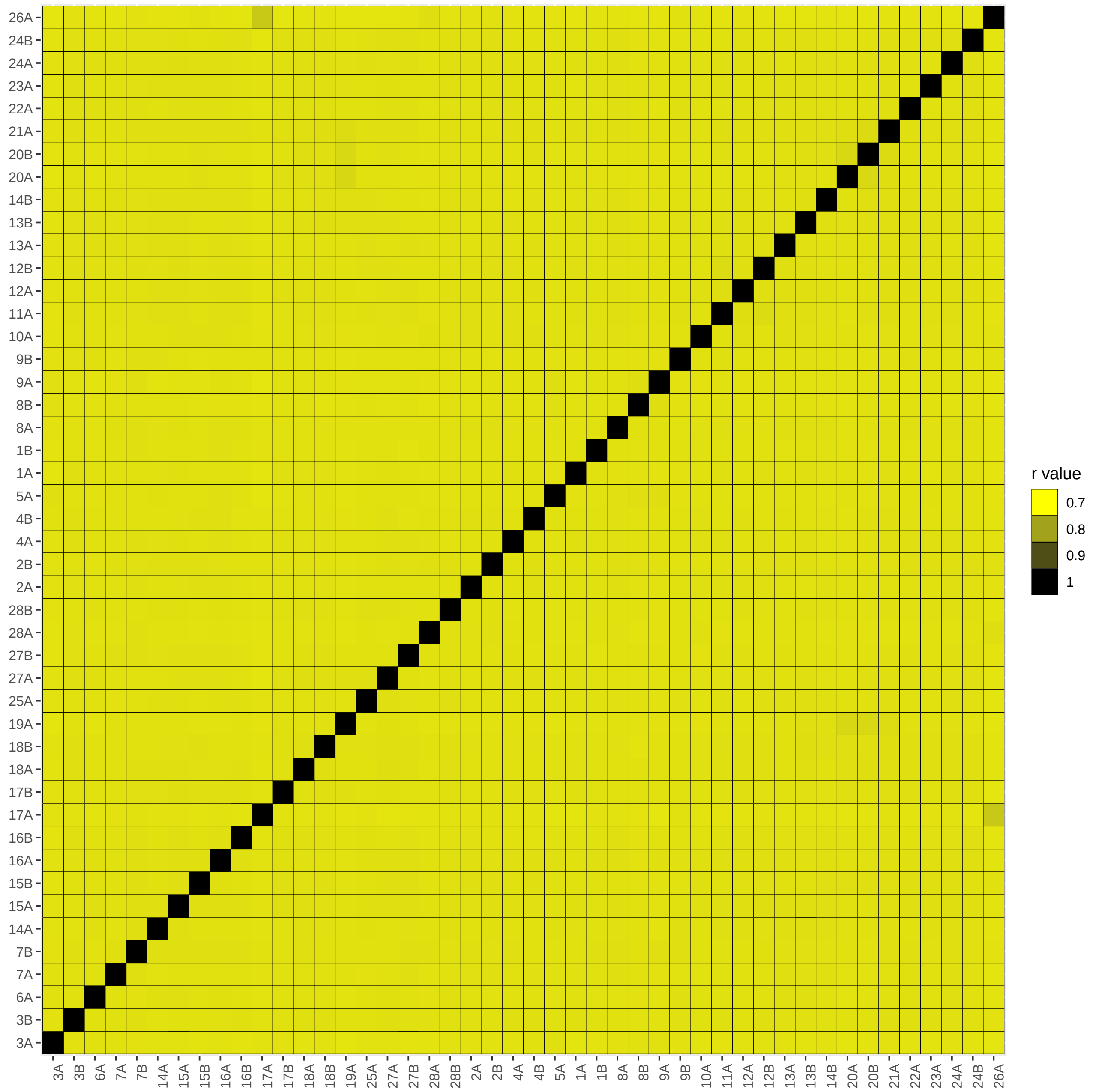


**Supplementary Figure S1. Identity-by-state analysis for wild-caught bumblebees.**

As genealogical information on our wild-collected bumblebees is unavailable, we performed an identity-by-state analysis using genome-wide polymorphic sites. This shows no evidence of substructure among the 46 bumblebees collected, indicating they originate from a single population. Heatmap shows the percentage of shared SNPs (r-value) calculated between two individual bumblebees. Sample names for each bumblebee are provided on the x and y axes.

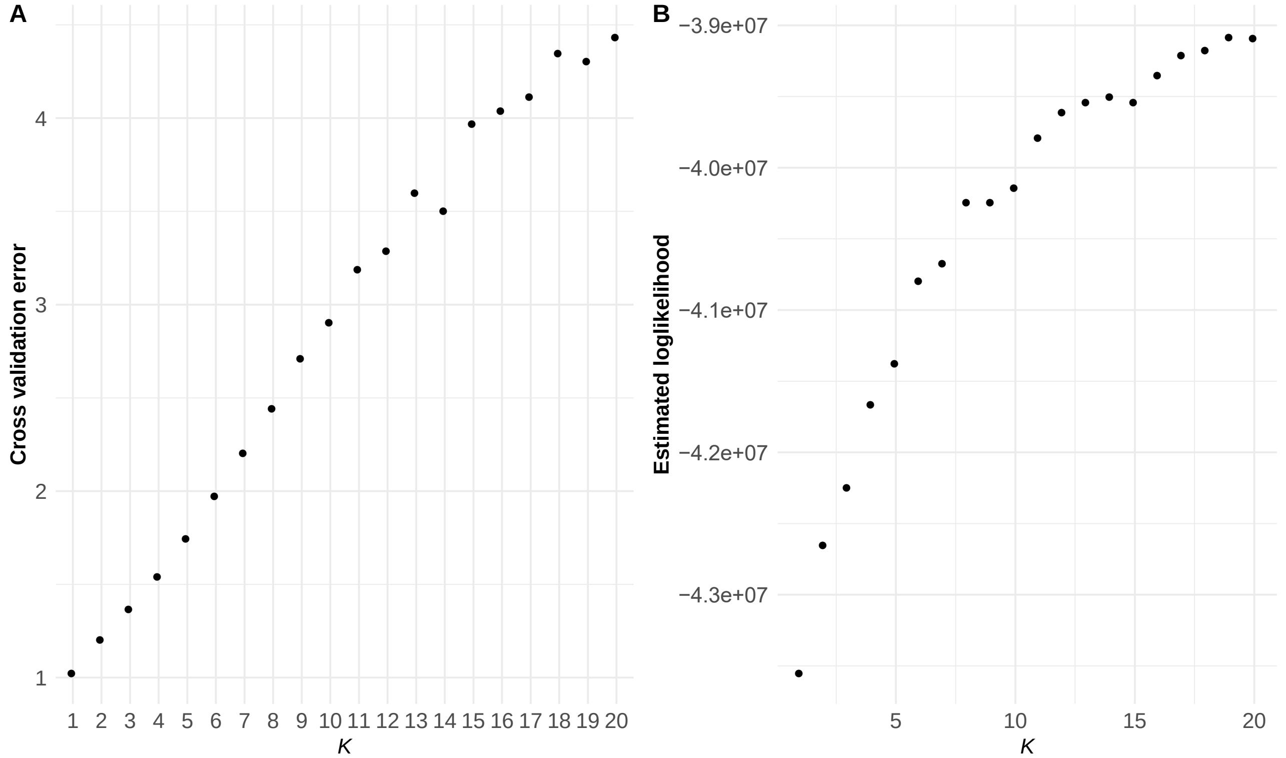


**Supplementary Figure S2. Weak population substructure within British *B. terrestris*.**

Scatterplot displaying estimates for the number of predicted populations in the current dataset as measured by ADMIXTURE: (A) cross-validation error rate and (B) estimated log-likelihood for each predicted *K* (1-20). The lowest cross-validation error rate within the current dataset was for *K*=1, suggesting that this species is a panmictic population on the island of Great Britain.

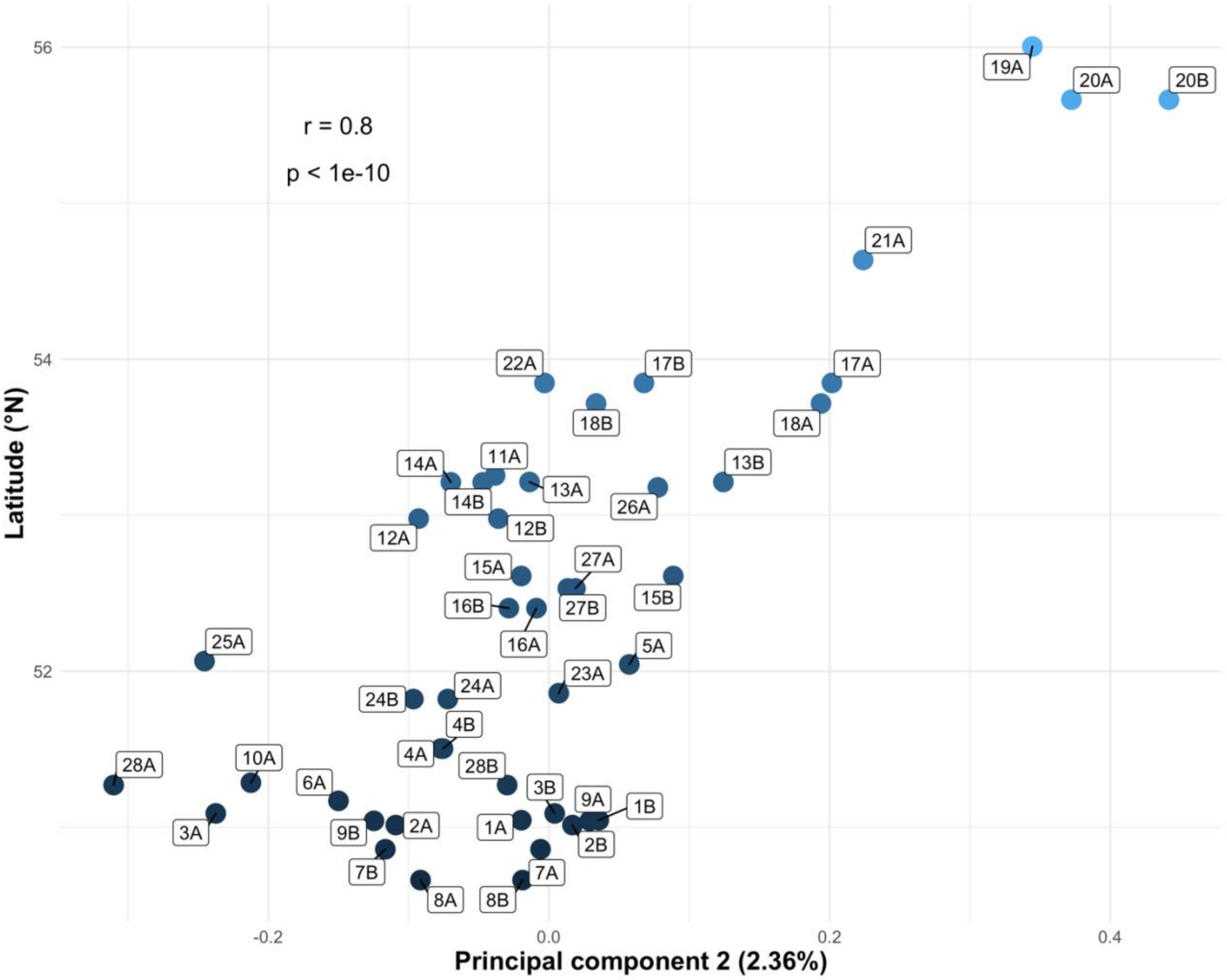


**Supplementary Figure S3. Correlation between genotype frequencies and latitude.**

Scatterplot displaying significant positive correlation (Pearson's r=0.8, $p<10^{-10}$) between latitude (°N) of site where bumblebees were collected and principal component 2 from a principal component analysis performed on genome-wide SNPs. Each dot represents an individual bumblebee male with the color representative of the latitude of the collection site. Each label contains a number (i.e., 1-28) and a letter ("A" or "B") with the number indicating a unique collection site while the letter indicates the first ("A") and second male ("B") collected from a particular site.

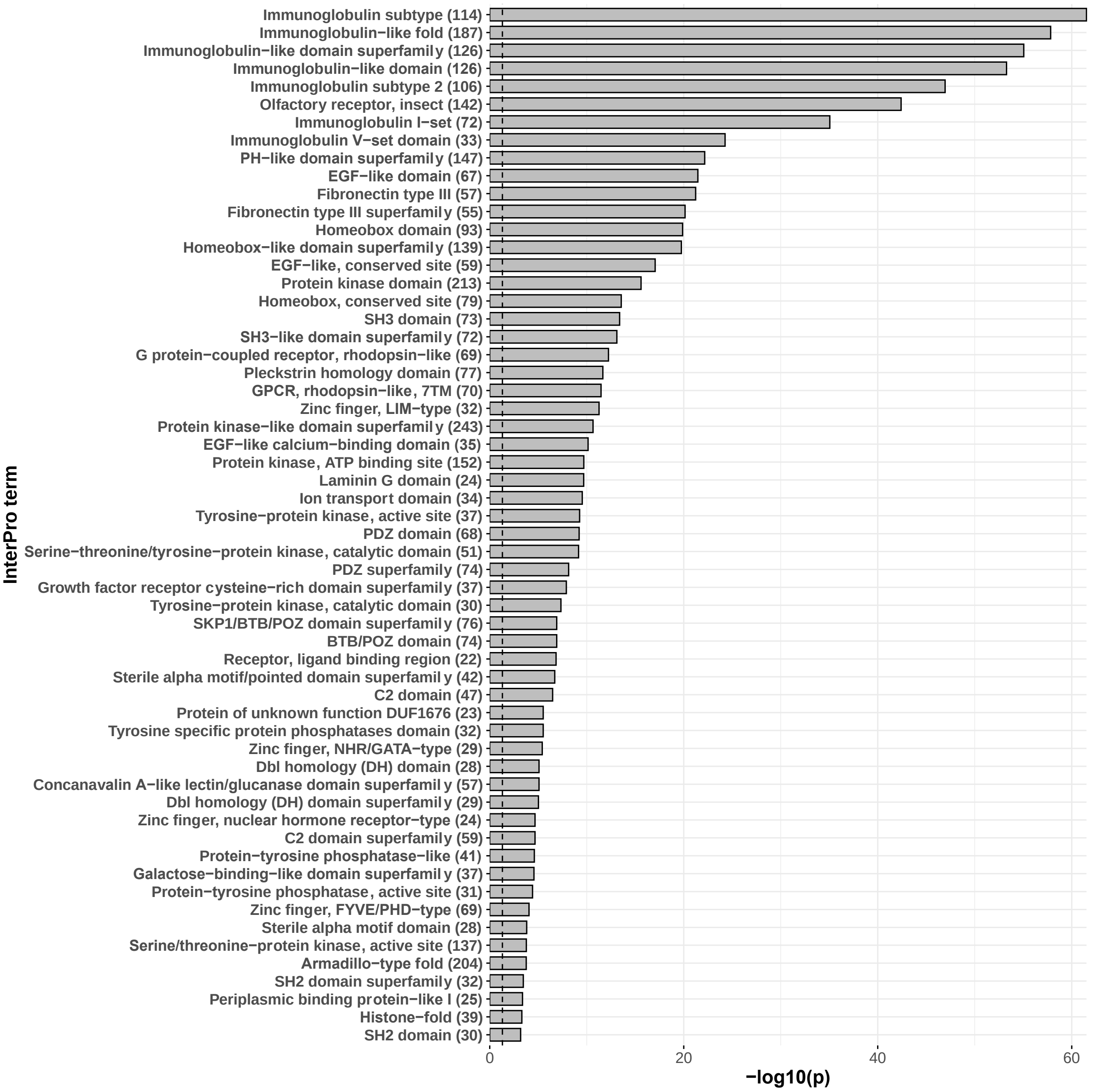


**Supplementary Figure S4. Enriched functional domains in bumblebee genes under recent selection.**

Bar plot displaying the -log10 transformed Bonferroni adjusted *p* values for InterPro domains enriched in genes with high |$nS_L$| scores. For each significantly enriched InterPro functional domain, we show its description and the total number of annotated terms in the *B. terrestris* proteome.

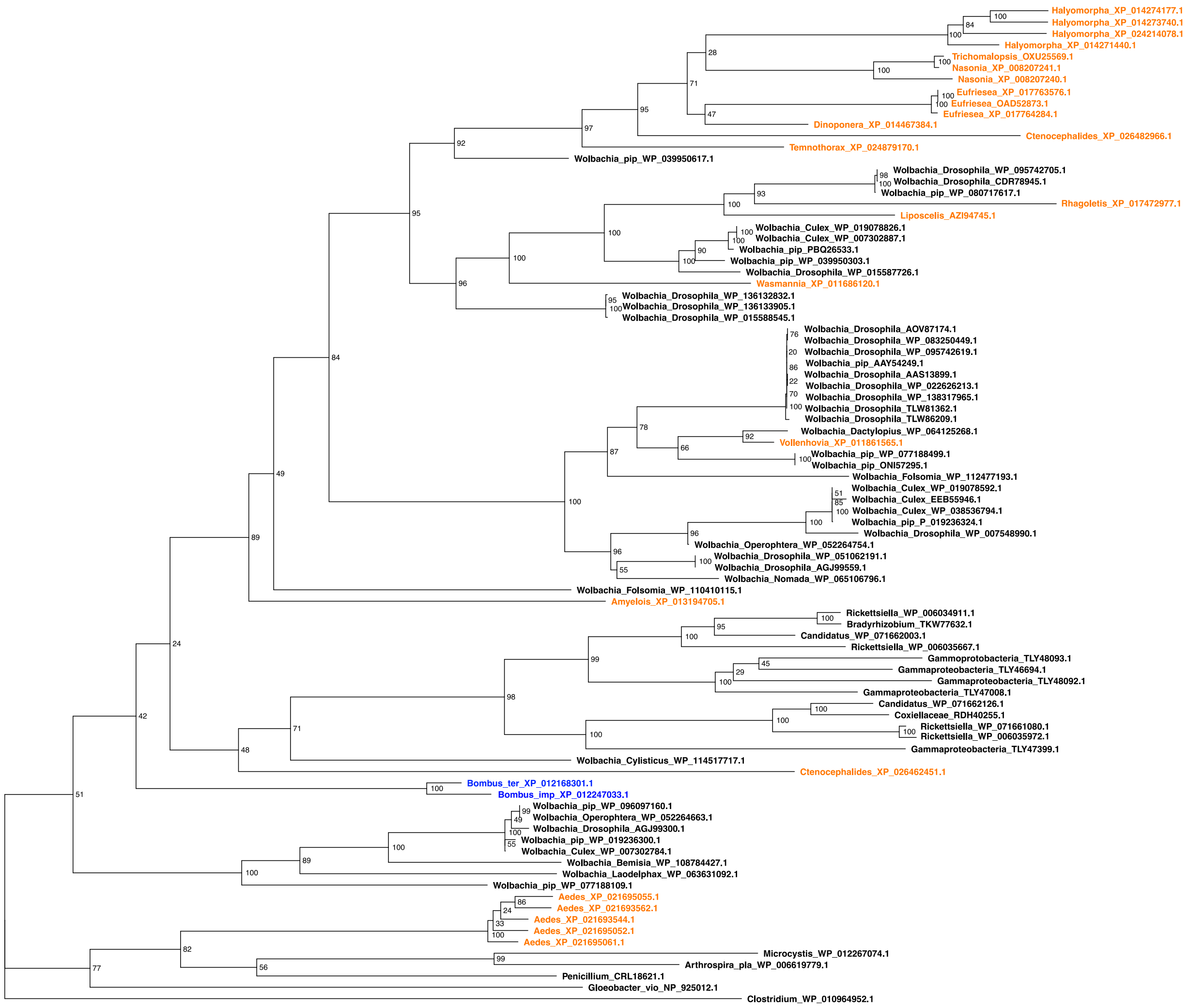


**Supplementary Figure S5. Phylogeny of LOC105666162, the gene horizontally transferred from *Wolbachia* into *Bombus* and similar sequences from other organisms.**

LOC105666162 in *Bombus terrestris* and its homolog in *B. impatiens* (in blue), are nested among sequences from prokaryotic genomes (black), with some other sequences annotated as being from eukaryotes (orange). Bootstrap values are provided for each node.

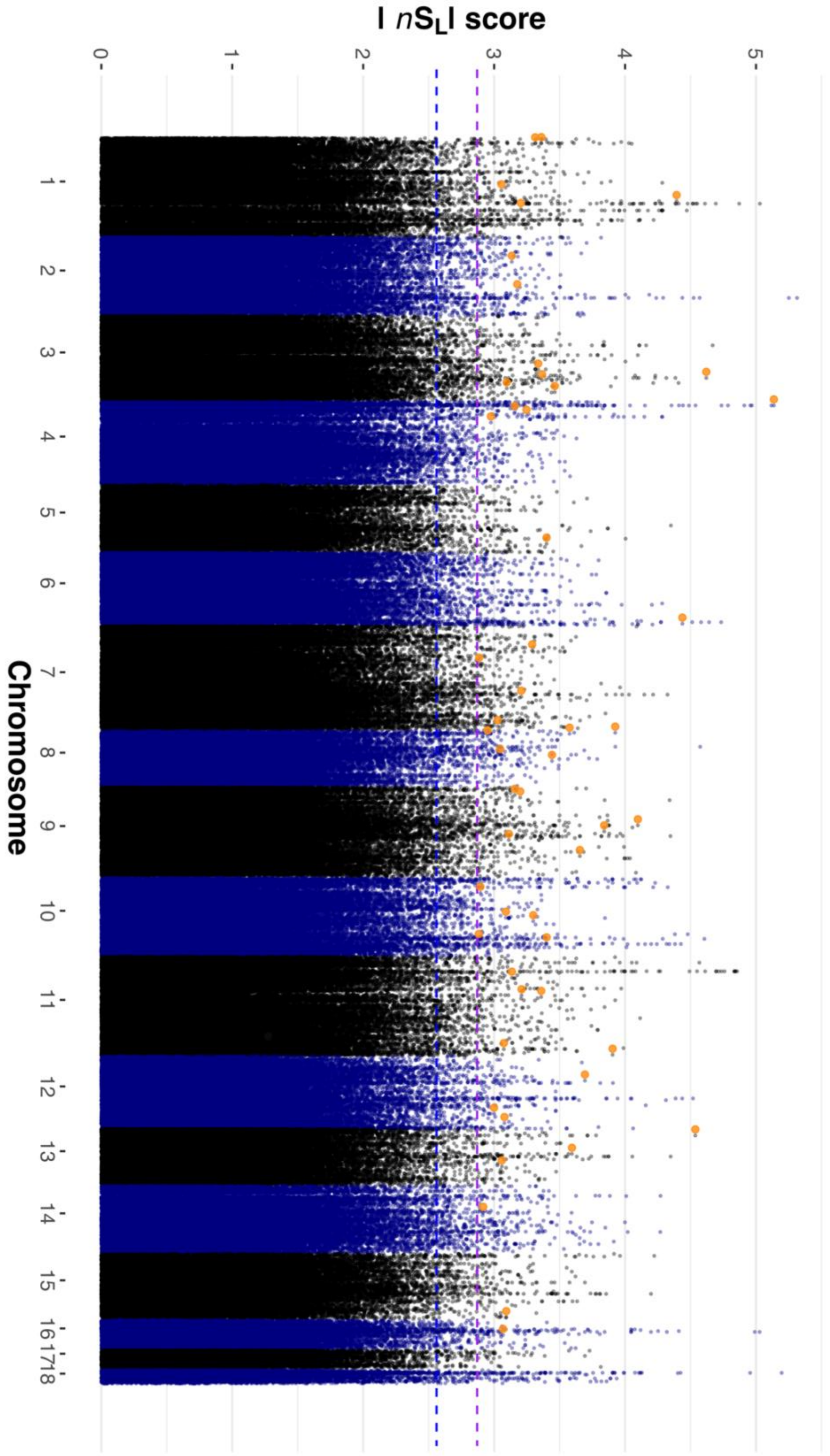


**Supplementary Figure S6. Putative pesticide-response genes under selection in the bumblebee genome.**

Genomic location and $|nS_L|$ score of all SNPs. For genes with putative roles in pesticide response, the highest $|nS_L|$ score is highlighted in orange. Blue and purple horizontal dashed lines respectively indicate the 1st percentile of overall $|nS_L|$ scores and 10th percentile of genic $|nS_L|$ scores.

#### Legends for Supplementary Tables 1 to 10

**Supplementary Table 1. Sample information for wild-caught bumblebees**
Sample information for male bumblebees collected across the island of Great Britain. For each bumblebee, we provide sample ID, collection site ID, latitude and longitude coordinates for collection site, the tissue(s) used for DNA extraction, the sequencing platform, as well as the number of sequencing runs, resulting numbers of paired-end reads generated per run, and overall predicted genome coverage.

**Supplementary Table 2. $nS_L$ analysis to identify bumblebees genes with signatures of recent positive selection**
The statistic $|nS_L|$ was calculated for each high quality SNP identified in the mainland British population with the SNP with highest $|nS_L|$ score per gene reported for genes found on one of the 18 chromosome-level scaffolds of the bumblebee genome reference assembly. For each gene, the table shows locus ID (NCBI gene symbol), top $|nS_L|$ score for that gene, the SnpEff functional annotation of the SNP with highest $|nS_L|$ score and gene description.

**Supplementary Table 3. Gene Ontology terms enriched in genes under recent positive selection**
Gene Ontology terms enriched in genes with signatures of recent positive selection. For each enriched term, the Gene Ontology identifier, term description, the number of genes annotated with that respective term, as well as the observed and expected values for each term and associated Bonferroni-adjusted p value ($p < 0.05$) are provided.

**Supplementary Table 4. Enrichment of InterPro domains in bumblebee genes with signatures of recent positive selection**
Enrichment of functional protein domains in bumblebee genes with signatures of recent positive selection. For each enriched functional protein domain, the InterPro domain ID, domain description, the number of predicted proteins annotated with IPR terms and measure of significance (Wilcoxon test; Bonferroni-adjusted $p < 0.05$) are shown.

**Supplementary Table 5. BLAST results of a protein coded for by an ancestral horizontal gene transfer event**
Results for the longest protein isoform (XP_012168301) of LOC105666162 BLASTp searches against the NCBI non-redundant (nr) database using the full predicted amino acid sequence, as well as individual searches for four subsections of unequal lengths.

**Supplementary Table 6. Gene expression across bumblebee life stages, castes and sexes**
Gene-level counts in transcripts per million for genes expressed across life stages, castes and sexes for two publicly available datasets. For each gene, NCBI RefSeq gene symbol is provided, while for each sample, individual sample description and SRA accession code is provided.

**Supplementary Table 7: Genes within conserved regions of low diversity on chromosome one**
Gene description and genomic coordinates for *Bombus terrestris* genes located in the region of low nucleotide diversity on chromosome one. We also provide the genomic coordinates of orthologs in the honeybee (*Apis mellifera*) genome assembly, orthology type, percentage similarity between orthologs and ortholog confidence score.

**Supplementary Table 8. Putative insecticide resistance genes under positive selection**
Top $|nS_L|$ score for genes belonging to gene families previously associated with insecticide resistance including targets of insecticides, such as nicotinic acetylcholine receptors (nAChRs), as well as genes involved in xenobiotic detoxification, such as cytochrome P450s, carboxylesterases and glutathione S-transferases.

**Supplementary Table 9. Putative pesticide response genes under positive selection**
Top $|nS_L|$ score for genes previously identified as differentially expressed or alternatively spliced in *Bombus terrestris* castes in response to neonicotinoid exposure.

**Supplementary Table 10. Copy number variation in *Bombus terrestris***
Raw median and normalized read depths for each sample for duplicated sites identified across a mainland British population of *Bombus terrestris*. For each putative duplicated site, the genomic position, CNV length (bp) and names (NCBI RefSeq gene symbol) of overlapping genes (if applicable) are provided.
